# *Prototheca zopfii* Genotype II induces mitochondrial apoptosis in models of bovine mastitis

**DOI:** 10.1101/562983

**Authors:** Muhammad Shahid, Eduardo R. Cobo, Liben Chen, Paloma A. Cavalcante, Herman W. Barkema, Jian Gao, Siyu Xu, Yang Liu, Cameron G. Knight, John P. Kastelic, Bo Han

## Abstract

*Prototheca zopfii* is an alga increasingly isolated from bovine mastitis. Of the two genotypes of *P. zopfii* (genotype I and II (GT-I and II)), *P. zopfii* GT-II is the genotype associated with acute mastitis and decreased milk production by unknown mechanisms. The objective was to determine inflammatory and apoptotic roles of *P. zopfii* GT-II in cultured mammary epithelial cells (from cattle and mice) and murine macrophages and using a murine model of mastitis. *Prototheca zopfii* GT-II (but not GT-I) invaded bovine and murine mammary epithelial cells (MECs) and induced apoptosis, as determined by the terminal deoxynucleotidyl transferase mediated deoxyuridine triphosphate nick end labeling assay. This *P. zopfii* GT-II driven apoptosis corresponded to mitochondrial pathways; mitochondrial transmembrane resistance (ΔΨm) was altered and modulation of mitochondrion-mediated apoptosis regulating genes changed (increased transcriptional *Bax*, cytochrome-c and *Apaf-1* and downregulated *Bcl-2*), whereas caspase-9 and -3 expression increased. Apoptotic effects by *P. zopfii* GT-II were more pronounced in macrophages compared to MECs. In a murine mammary infection model, *P. zopfii* GT-II replicated in the mammary gland and caused severe inflammation with infiltration of macrophages and neutrophils and upregulation of pro-inflammatory genes (*TNF-α*, *IL-1β* and *Cxcl-1*) and also apoptosis of epithelial cells. Thus, we concluded *P. zopfii* GT-II is a mastitis-causing pathogen that triggers severe inflammation and also mitochondrial apoptosis.

**Author summary:** Bovine mastitis (inflammation of the udder) reduces milk production and quality, causing huge economic losses in the dairy industry worldwide. Although the alga *Prototheca zopfii* is a major cause of mastitis in dairy cows, mechanisms by which it damages mammary tissues are not well known. Here, we used cell cultures and a mouse model of mastitis to determine how *Prototheca* caused inflammation and cell death in mammary tissues. *Prototheca* invaded mammary gland cells, from cattle and mice, as well as macrophages (white cells that take up and kill pathogens) and caused cell death by interfering with mitochondria. Furthermore, *Prototheca* causes severe inflammation and tissue damage when injected into the mammary glands of mice. Although there are two genotypes of *P. zopfii*, only genotype II causes tissue damage, whereas gentotype I, common in farm environments, does not damage mammary tissues. Since *P. zopfii* is an alga and not a bacterium, antibiotic treatments, frequently used to treat mastitis in cattle, are not effective against this organism. Understanding how *P. zopfii* damages mammary tissue and causes mastitis is important new knowledge to promote future development of evidence-based approaches to prevent and treat mammary gland infections with this organism.

## Introduction

Bovine mastitis (inflammation of the udder), caused by infection with pathogenic microorganisms and destruction of milk-synthesizing tissues [1], reduces milk production and quality and is an important financial threat to the dairy industry [2]. *Prototheca zopfii*, a chlorophyllous alga (family *Chlorellaceae*) unable to synthesize chlorophyll and with heterotrophic modes of nutrition [3,4], is a major cause of mastitis in dairy cows [5,6]. Bovine protothecal mastitis can be clinical or subclinical. In clinical cases, symptoms include fever (up to 40 °C), pain, mammary edema, anorexia and reluctance to move [7]. Subclinical protothecal mastitis is associated with increased number of leukocytes in the udder and milk, and can be manifested by slight pain along with loss of appetite [7]. Bovine protothecal mastitis decreases milk production and elevates somatic cell count in milk, especially macrophages, often resulting in culling [7]. Reported bovine *Prototheca zopfii* mastitis occurrence ranges from 7.5 to 16.3% [8,9]; however, these reports are predominantly from outbreaks. Although a large proportion (up to 81%) of dairy herds are infected, this pathogen affects a limited proportion of cows (<10%) [10,12]. Cows are often infected intramammarily with *P. zopfii* following teat trauma during mechanical milking [13] and contamination of the teat orifice with damp organic material [7,14]. Single *Prototheca zopfii* endospores or sporangiospores contact mammary gland epithelial cells, which are first responders, sensing their presence and initiating an inflammatory immune response. After breaching epithelial defenses, *Prototheca zopfii* may also invade macrophages of the mammary gland alveolar lumen and interstitium [15], making *Prototheca zopfii* less accessible to antibiotics and diagnostic methods [16].

Two genotypes of *Prototheca zopfii,* genotype I (GT-I) and genotype II (GT-II) have been isolated and identified from bovine milk [17]. Genotype I is predominantly isolated from environmental samples, whereas GT-II is isolated from milk samples and has been reported as the causative pathogen of bovine mastitis [11,18,19]. *Prototheca zopfii* GT-II induced oxidative stress and apoptotic death in cultured bovine mammary epithelial cells (bMECs) [20,21], but pathogenesis of protothecal mastitis remains elusive. Thus, we aim to determine inflammatory and apoptotic roles of *Prototheca zopfii* GT-II in cultured mammary epithelial cells (from cattle and mice) and murine macrophages and using a murine model of mastitis.

## Materials and methods

### Statement of ethics

The current study was conducted in accordance with ethical guidelines and regulations regarding laboratory animal care and use, as described in the “Guide to the Care and Use of Experimental Animals” from the Canadian Council on Animal Care (https://www.ccac.ca/Documents/Standards/Guidelines/Experimental_Animals_Vol1.pdf). Animal use was reviewed and approved by the Animal Care Committee of the University of Calgary, Calgary, AB, Canada (protocol number AC16-0061).

### Prototheca zopfii culture

*Prototheca zopfii* GT-II isolates were collected from milk samples of dairy cows with clinical mastitis, whereas *P. zopfii* GT-I isolates were predominantly cultured from environmental samples in China, and cultured and stored at College of Veterinary Medicine, China Agricultural University, Beijing, China [22]. Prior to each experiment, fresh *P. zopfii* GT-I and -II were cultured on Sabouraud dextrose agar (SDA; Sigma, Shanghai, China) for up to 48 h at 37°C and single colonies incubated in Sabouraud dextrose broth (SDB; Sigma) at same conditions for up to 72 h [22].

### Mouse protothecal mastitis model

C57BL/6 lactating female mice (6–8 wk old; 10-14 d after parturition) were housed in specific pathogen-free facilities at the University of Calgary with *ad libitum* access to food and water. Mice were inoculated intramammarily with either *P. zopfii* GT-II (50 µL containing 1 × 10^5^ CFU/mL) or an equal volume of phosphate buffered saline (PBS) (control) in the left fourth and right fourth (L4 and R4) mammary glands. Mice were euthanized 4 d post inoculation (dpi) to collect mammary tissue samples. Tissues were mixed into TRIzol (Invitrogen, Carlsbad, CA, USA) and later homogenized for quantitative PCR (qPCR) or fixed in 10% formalin solution, embedded in paraffin wax, sectioned with a microtome (5 µm) and stained with hematoxylin and eosin (H&E; Sigma, USA) for histological examination [23], and Periodic Acid-Schiff (PAS; Sigma, USA) and Grocott-Gomori’s methenamine silver stain (GMS) as a screen for fungal organisms).

### Identification of macrophages and neutrophils in murine mammary gland

Fixed murine mammary gland tissue sections were deparaffinized, dehydrated and permeabilized with PBS/Triton X-100 (0.25%, v/v) (PBS-T) buffer containing 1% donkey serum (Cat # 017-000-121) at room temperature for 10 min. Slides were blocked with PBS-T containing 10% (v/v) donkey serum and 1% (v/v) bovine serum albumin (BSA) (Sigma, USA) for 120 min at room temperature. After washing with PBS, sections were incubated with primary antibodies against murine F4/80 (macrophages) (Cat # 4316835, BD Pharmingen™, US) and Ly-6G (neutrophils) antigens (Cat# 127609, Biolegend, US) (1:1,000 in PBS-T plus 1% BSA) for 16 h at 4°C. Following washing with PBS-T, slides were incubated with secondary antibodies (488-conjugated Affinipure Goat anti-Rat IgG, Cat# 135205, Jackson Immune Research, UK) (1:1,000 in PBS-T plus 1% BSA) at room temperature for 60 min and washed again with PBS-T and then incubated with DAPI (4’, 6-diamidino-2-phenylindole) (Invitrogen) at room temperature for 20 min. Slides were examined with an immunofluorescence microscope (ZEISS Axio Imager M2, Carl Zeiss AG, Jena, Thuringia, Germany).

### Epithelial cell and macrophage culture

A bMEC line isolated from a cow (MAC-T) (Shanghai Jingma Biological Technology Co., Ltd. China), murine macrophages derived from mouse BALB/c monocytes (J.774, provided by Dr. Eduardo R. Cobo, University of Calgary) and a murine mammary epithelial cells line (mMECs; HC11, provided by Dr. Eduardo R. Cobo, University of Calgary) were used. The bMECs and murine macrophages were cultured in HyClone™ DMEM/F12 medium (Thermo Fisher Scientific, South Logan, NH, USA) along with 10% fetal bovine serum (FBS; Thermo Fisher Scientific), penicillin (100 U/mL; Thermo Fisher Scientific) and streptomycin (100 U/mL; Thermo Fisher Scientific) in cell culture plates (Corning Inc., Corning, NY, USA). The mMECs were cultured in RPMI (Thermo Fisher Scientific) medium along with 10% fetal bovine serum (FBS; Thermo Fisher Scientific), penicillin (100 U/mL; HyClone®, USA) and streptomycin (100 U/mL; Thermo Fisher Scientific). For experimental challenges, bMECs and macrophages (bovine and murine) were challenged with *P. zopfii* GT-I and GT-II suspended in DMEM/F12 to 5 × 10^5^ and 1 × 10^5^ CFU/mL, respectively, for up to 24 h at 37°C with 5% CO_2_.

### *P. zopfii* cell internalization assay

Murine macrophages and bMECs were infected with *P. zopfii* for up to 8 h, washed with PBS (pH 7.4) and incubated for 2 h with gentamycin (200 μg/mL) to eliminate extracellular *P. zopfii*. Cells were washed with PBS to eliminate non-adherent bacteria and then lysed by 0.5% Triton X-100 (v/v) to determine CFU by 10-fold serial dilution [24]. Further confirmation of phagocytic activity of macrophages was conducted by actin inhibition (cytochalasin D; C8273, Sigma, USA; 1 h) before inoculation.

### Transmission electron microscopy (TEM)

Bovine MECs infected with *P. zopfii* GT-I and -II were washed with PBS (pH 7.2), fixed with 2% glutaraldehyde and 1% paraformaldehyde (pH 7.2; Sinopharm Chemical Reagent Co., Shanghai, China) and processed for TEM [21].

### Mitochondrial damage assay

After infection with *P. zopfii,* GT-I and –II, bovine MECs were collected to assess changes in mitochondrial membrane potential (ΔΨm) as determined by the presence of JC-1 (Cat# M8650, Solarbio, Beijing, China) using flow cytometry and immunofluorescence microscopy. JC-1 is a dual-emission potential-sensitive probe that forms red-fluorescent aggregates in healthy mitochondria, but becomes a green-fluorescent monomer after membrane potential collapses.

### Transcriptional gene expression of inflammatory and apoptotic genes

Total RNA was extracted from bMECs, mMECs and murine macrophages with TRIzol reagent (Invitrogen) and converted to cDNA (RevertAid First Strand cDNA synthesis kit, Thermo Scientific). The resulting RNA and cDNA quality was evaluated by the absorbance ratio (A260/A280 ratio) (NanoVue Spectrophotometer, GE Healthcare Bio-Sciences, Little Chalfont, Buckinghamshire, UK) [25], which was corrected to be ∼1.8–2.0 for an individual sample. Amplification of mRNA genes for *TNF-α*, *IL-1β*, *IL-8/Cxcl-1*, *Bcl-2*, *Bax*, *Apaf-1*, cytochrome-c, caspase-9 and caspase-3 was done using a CFX-96 real-time PCR system (BioRad, Hercules, CA, USA). The reaction mixture for each sample carried 2 µL of cDNA, 1X SsoAdvanced Universal SYBR Green Supermix (BioRad) and 0.5 μM of each specific primer, in a 10 μL final volume. Relative primers for bovine and murine genes are shown (Tables 1 and 2, respectively). Reaction mixtures were incubated at 95°C for 5 min, followed by denaturation for 5 s at 95°C and combined annealing/extension for 10 s at 60°C (total of 40 cycles). All treatments were examined in duplicate in three independent experiments. The values of target mRNA were corrected relative to the normalizer *GAPDH*. Data were assessed using the 2−ΔΔCT method [25] and results presented as mean fold change of target mRNA levels in infected groups versus an uninfected control group [25].

**Table 1.**
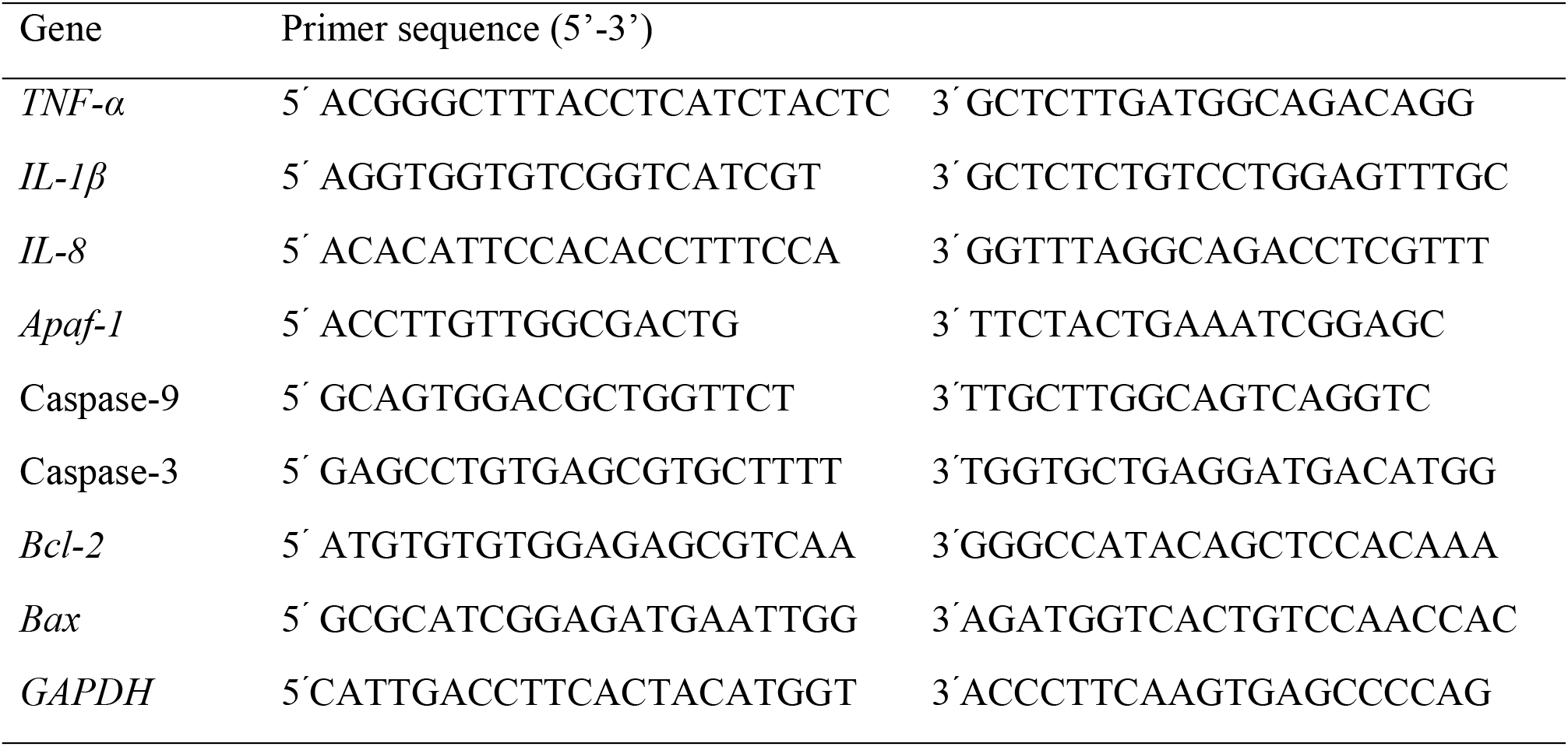
Primer sequences of qPCR for bovine genes.

**Table 2.**
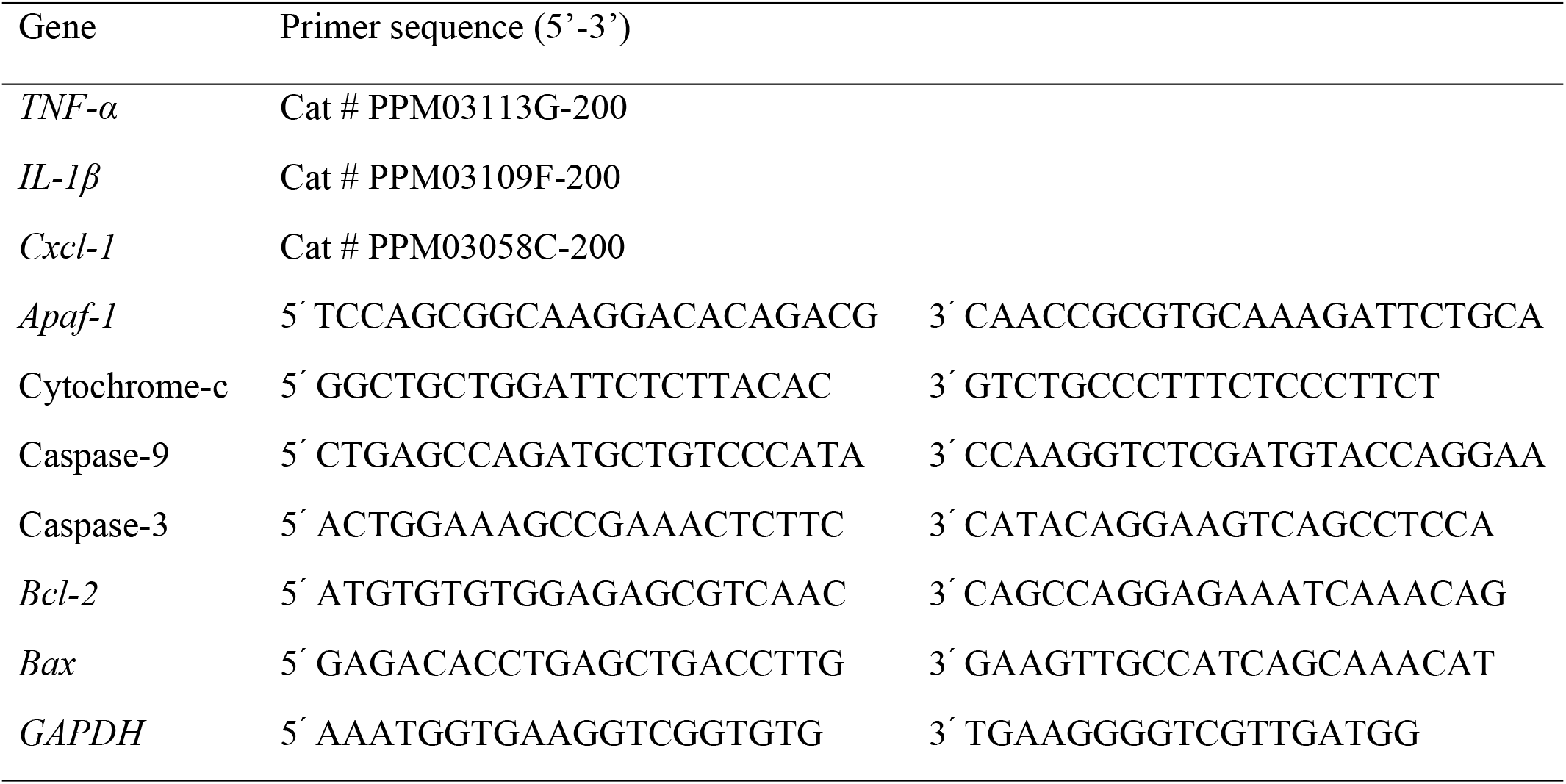
Primer sequences of qPCR for murine genes.

### TUNEL apoptosis staining

Apoptosis of bMECs, mMECs, murine macrophages and mouse mammary gland after *P. zopfii* GT-II inoculation was assessed by *in situ* TUNEL staining (S7165 ApopTaq apoptosis detection kit, MilliporeSigma, Haverhill, MA, USA). Apoptotic indices were calculated as positive stained apoptotic cells per field, using five fields per sample at 400 × magnification.

### Protein determination of apoptotic cytochrome-c, caspase-9, and caspase-3

Proteins from bMECs or homogenized murine mammary tissue were size-separated by SDS-PAGE and transferred to Immobilon-P polyvinylidene difluoride (PVDF) membrane (0.45 µm) (Millipore Sigma, Gillingham, Dorset, UK). Membrane was blocked with 5% skim milk in TBS-T (150 mM NaCl, 10 mM Tris base, 0.05% Tween 20, pH 7.4) at room temperature for 120 min and then incubated overnight at 4°C with primary antibodies for caspase-9 (Cat # ab69514, Abcam USA), caspase-3 (Cat # ab90437, Abcam USA), cytochrome-c (Cat # ab110325, Abcam USA) and housekeeping β-tubulin (Cell Signaling Technology, Danvers, MA, USA). The membrane was rinsed with TBS-T and incubated with HRP-labeled secondary goat anti-rabbit IgG (ZRA03, Biotech, China) or goat anti-mouse IgG (ZM03, Biotech, China) at 37°C for 60 min. Signals were detected using enhanced chemiluminescence (Cat # PE0010, Solarbio Life Sciences, Beijing, China).

### Statistical analyses

Data were analyzed in triplicate for reproducibility and were expressed as mean ± standard deviation (SD). Data of the infected and uninfected groups were analyzed using paired Student’s *t-*test with 95% confidence interval. Data were further analyzed by ANOVA and *post hoc* tests using SPSS 20.0 (International Business Machines Corporation, Armonk, NY, USA). For all analyses, *P* < 0.05 was considered significant.

## Results

### *P. zopfii* GT-II induced mastitis and apoptosis in a mouse model

To investigate the causative effect of *P. zopfii* GT-II in protothecal mastitis, lactating mice were intramammarily challenged with *P. zopfii* GT-II isolated from a bovine clinical mastitis case. Round to oval sporangia with regular internal divisions compatible with *P. zopfii* were observed in the mammary gland of lactating mice at 4 dpi as detected by PAS and GMS staining (Fig 1A). *Prototheca zopfii* GT-II replicated in the murine mammary gland as it was recovered by culture in greater amounts at 4 dpi compared to the initial inoculum (mean 3.4 × 10^7^ CFU/g tissue).

**Fig 1.**
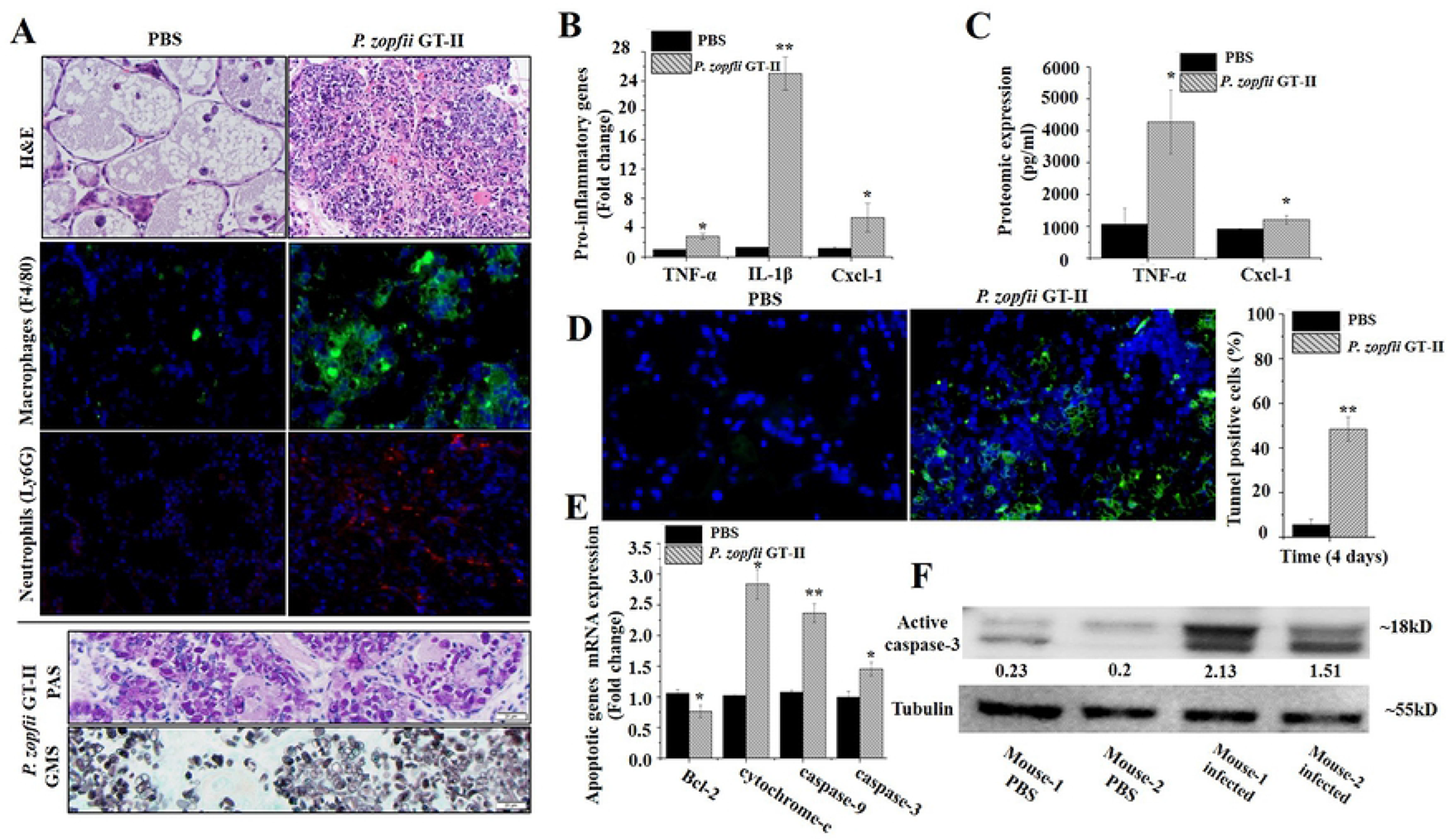
*Prototheca zopfii* genotype II-induced microscopic changes in the mammary gland of mice. (**A**) H&E, PAS and GMS staining and immune detection of macrophages (F4/80) and neutrophils (Ly6 G) in PBS control and *P. zopfii*-infected mammary tissues at 4 d post infection. Note infiltration of macrophages and neutrophils and the presence of innumerable *P. zopfii* GT-II as detected by PAS and GMS staining. Bar = 20 µm. (**B**) Transcriptomic expression of genes of pro-inflammatory *TNF-α, IL-1β* and *Cxcl-1* after infection with *P. zopfii* GT-II. (**C**) ELISA titers of *TNF-α* and *Cxcl-1* in mammary tissue. (**D**) Quantitative detection of apoptotic cells in PBS control and *P. zopfii* GT-II infected mammary tissue in mouse mammary gland (green signal indicates TUNEL apoptotic cells). (**E**) Transcriptomic expression of *Bcl-2*, cytochrome-c, caspase-9, and caspase-3 in mouse mammary tissues infected with *P. zopfii* GT-II. (**F**) Proteomic expression of activated caspase-3 in mammary gland 4 d after *P. zopfii* GT-II infection. \**P* < 0.05, \*\**P* < 0.01.

*Prototheca zopfii* GT-II induced acute mastitis with infiltration of leukocytes throughout the parenchyma and within lumina of alveoli. *Prototheca zopfii* GT-II were present both free within alveolar lumina and throughout the interstitium of the mammary tissue (Fig 1A). Using immune detection, macrophages were demonstrated in the mammary interstitium and neutrophils diffusely distributed in *P. zopfii* GT-II-infected mice (Fig 1A). The presence of *P. zopfii* GT-II upregulated gene activity and protein production of pro-inflammatory *TNF-α, IL-1β* and *Cxcl-1* in mammary tissue at 4 dpi (Fig 1B-C).

Next, we determined whether intramammary infection with *P. zopfii* GT-II involved apoptosis and oxidative stress, as described in cultured bovine mammary epithelial cells (bMECs) [20,21]. Apoptotic cells were quantified at 4 dpi with *P. zopfii* GT-II (Fig 1D). Transcriptomic analysis demonstrated that mRNA expression of caspase-9 and caspase-3 genes regulating mitochondrion-mediated apoptosis was higher in *P. zopfii* GT-II infected mice (Fig 1E) with cleavage of caspase-3 protein (Fig 1F). Expression of *Bax* gene increased in mammary tissue after *P. zopfii* GT-II inoculation (Supplementary Fig 1A), whereas expression of *Bcl-2* decreased (Fig 1E). Expression of cytochrome-c released into the cytosol to trigger apoptosis (Fig 1E) and *Apaf-1* also increased in *P. zopfii* GT-II inoculated mice (Supplementary Fig 1B).

### *P. zopfii* GT-II-driven apoptosis occurred in both mammary epithelial cells and macrophages

Since mastitis is a process involving epithelial cells and leukocytes, we investigated contributions of single-cell components in the pathogenesis of *P. zopfii* GT-II mastitis and apoptotic responses. We used a murine MEC (HC11) with ability to produce milk proteins (beta-casein) in response to prolactin [26]. Infection with *P. zopfii* GT-II in MEC induced early *IL-1β, TNF-α* and *Cxcl-1* gene expression (after 2 hpi) (Fig 2A-C). Apoptotic cells appeared later (24 hpi; Fig 2D) with an increased transcriptional expression of hallmark apoptotic genes, *Bax*, *Apaf-1* (Supplementary Fig 1C-D), cytochrome-c, caspase-9 and -3 genes (Fig 2E-G). Expression of *Bcl-2* was reduced (Fig 2H).

**Fig 2.**
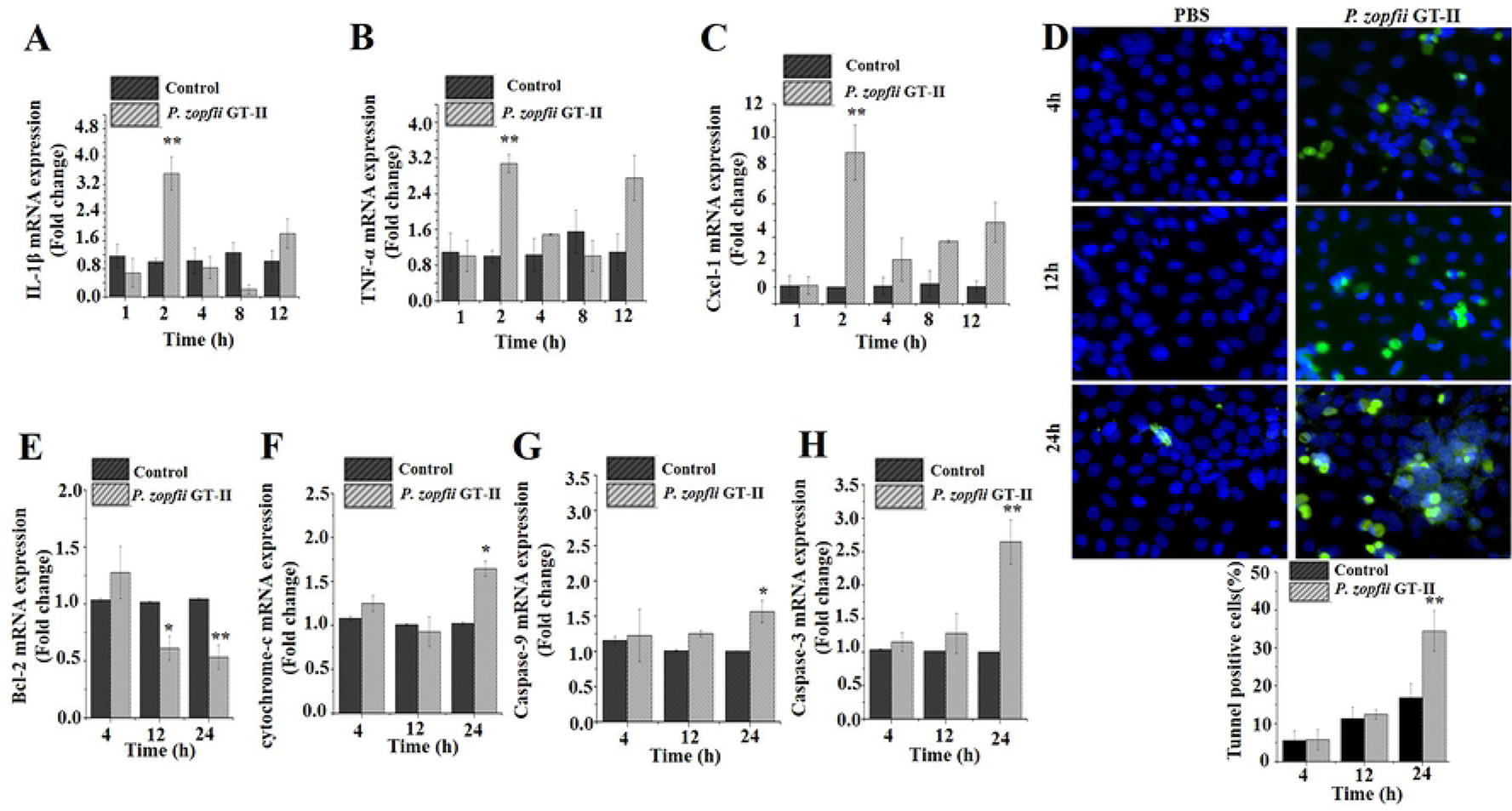
Murine MECs infected with *Prototheca zopfii genotype II* (GT-II). (**A, B and C**) Expression of mRNA level of *TNF-α, IL-1β*, and *Cxcl-1* in murine MECs was quantified after infection of *P. zopfii* GT-II infection. (**D**) Representative picture of TUNEL assay, plus quantitative analysis of TUNEL-positive apoptotic cells (20x). (**E, F, G and H**) mRNA expression of *Bcl-2*, caspase-9 and caspase-3, respectively, was quantified by qPCR and expressed as fold change relative to uninfected cells. Data are mean ± SD of three independent experiments. \**P* < 0.05, \*\**P* < 0.01.

To examine the role of macrophages, key in chronic mastitis [27], murine macrophages (J774) with phagocytic characteristics were challenged with *P. zopfii* GT-II. *Prototheca zopfii* GT-II internalized inside macrophages in a time-dependent fashion (up to 8 hpi; Fig 3A). This internalization seemed to be an active microbe process (*P. zopfii* dependent) rather than a phagocytic event, as actin inhibition in macrophages (by cytochalasin D) did not prevent *P. zopfii* GT-II internalization (Fig 3A). Infection of *P. zopfii* GT-II in macrophages upregulated mRNA expression of *IL-1β, TNF-α* and *Cxcl-1* (2 h; Fig 3B-D). In contrast, *TNF-α* expression decreased over time (Fig 3C). *Prototheca zopfii* GT-II induced apoptosis in macrophages as detected by TUNEL assay, with more cell death at 12 and 24 hpi (Fig 3E) and upregulated expression of *Bax, Apaf-1* (Supplementary Fig 1E-F), cytochrome-c, caspase-9, and caspase-3 genes, whereas *Bcl-2* expression decreased in a time-dependent manner (Fig 3F-I).

**Fig 3.**
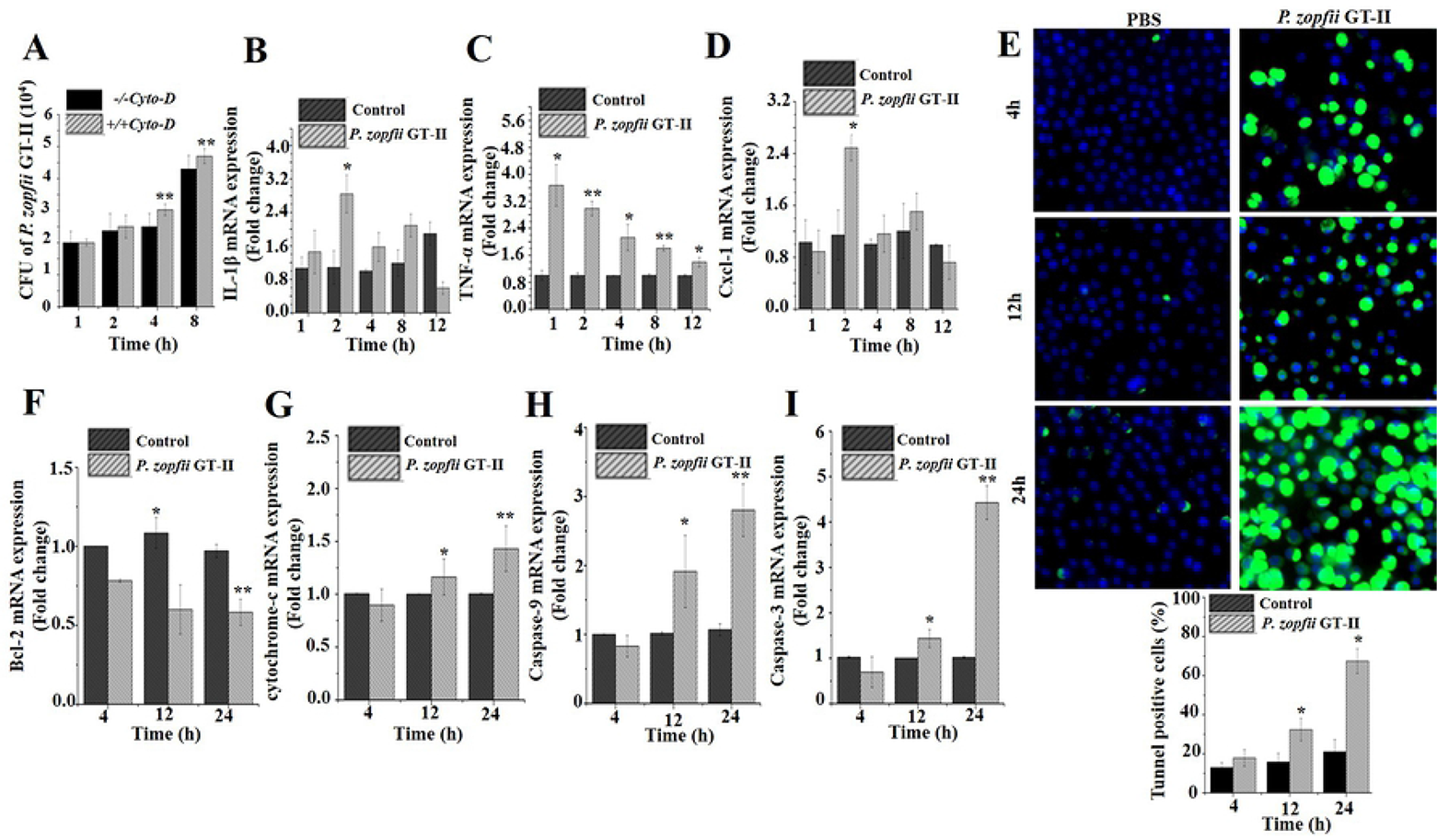
Murine macrophages infected with *P. zopfii* GT-II. (**A**) Internalization of *P. zopfii* GT-II in mouse macrophages in time-dependent manner, with and without cytochalasin D. (**B, C and D**) Level of cytokines (*TNF-α, IL-1β* and *Cxcl-1*) in murine macrophages. (**E**) TUNEL assay of mouse macrophages, quantitative analysis of apoptotic positive cells TUNEL positive apoptotic cells (20x). (**F, G, H and I**) Transcriptomic expression of *Bcl-2*, cytochrome-c, caspase-9, and caspase-3 at 4, 12 and 24 h after infection with *P. zopfii* GT-II in mouse macrophages on qPCR analysis and expressed as fold change relative to uninfected cells. Data are mean ± SD of three independent experiments. \**P* < 0.05, \*\**P* < 0.01.

### *P. zopfii* GT-II induced apoptosis in bovine mammary epithelial cells

To verify the apoptotic effects of *P. zopfii* in the target animal species (cattle), prototype bovine MECs with morphological and functional characteristics of normal mammary epithelial cells were challenged with *P. zopfii* GT-II and GT-I common commensals in farm environments (e.g., animal bedding, soil) [3]. *Prototheca zopfii* GT-I did not induce any apoptotic effects, but *P. zopfii* GT-II caused TUNEL-mediated apoptosis in a time-dependent manner (Fig 4A). This occurred rapidly, as *P. zopfii* GT-II were internalized by bMECs in the first 4 hpi, as confirmed by culture (Fig 4B) and TEM (Fig 4C). Apoptotic effects induced by *P. zopfii* GT-II were likely of mitochondrial origin, as mitochondrial transmembrane depolarization was detected by immunofluorescence and flow cytometry (12-24 hpi; Fig 4D-E).

**Fig 4.**
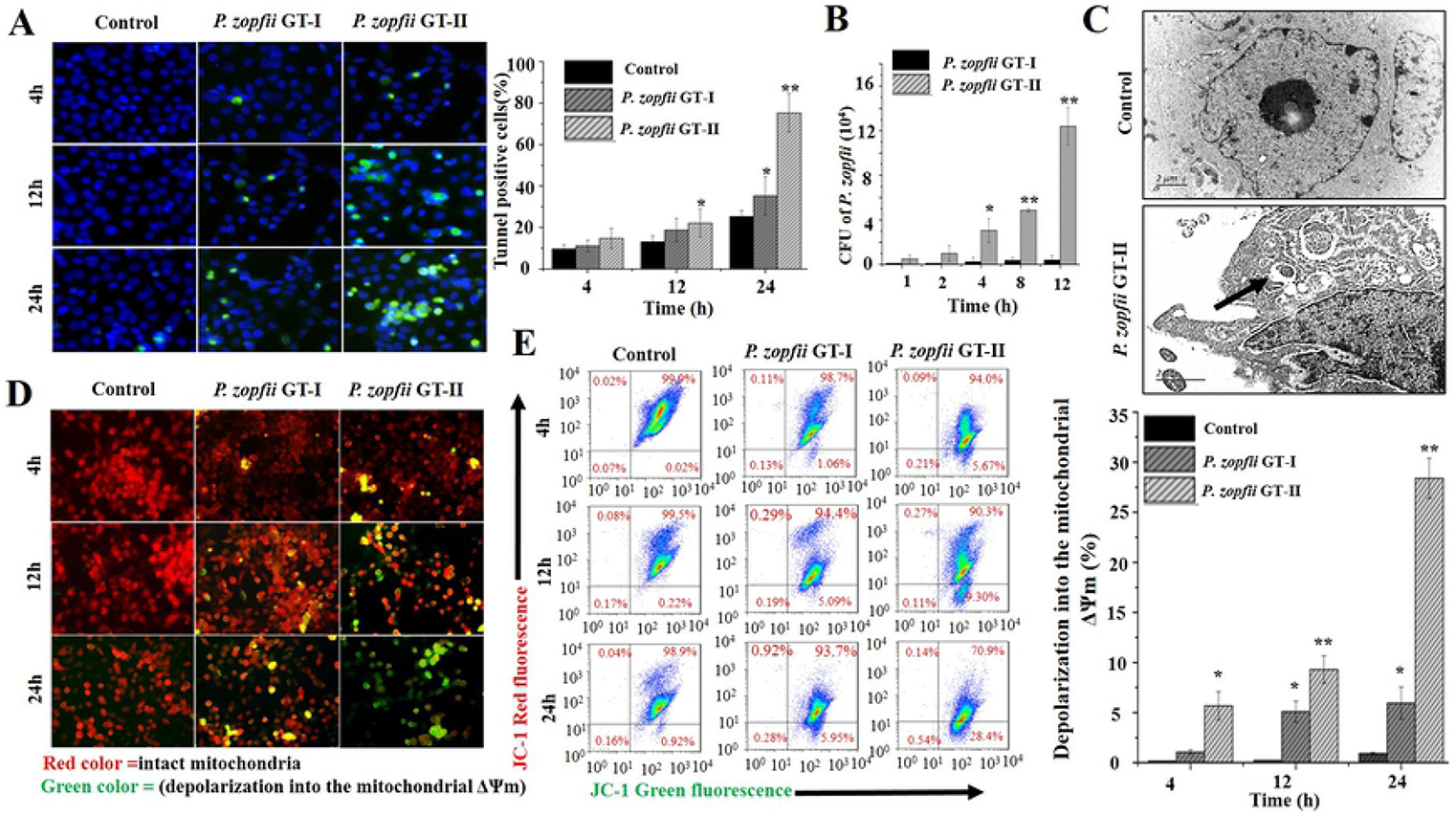
Bovine mammary epithelial cells (bMECs) *Prototheca zopfii genotype* (GT)-I and -II *in vitro* infection model. (**A**) Quantitative detection of apoptotic cells in *P. zopfii* GT-I and -II infected bovine mammary epithelial cells (green signal indicates TUNEL apoptotic cells). (**B**) Internalization of *P. zopfii* GT-II in bMECs was increased in a time-dependent fashion as compared to *P. zopfii* GT-I infection in bMECs, **(C)** Intracellular localization of *P. zopfii* GT-II in bMECs on transmission electron microscopy (black arrow). (**D**) Mitochondrial transmembrane potential (ΔΨm) assay of bMECs infected with *P. zopfii* using JC-1 staining (the compound 5,5’,6,6’-tetrachloro-1,1’,3,3’-tetraethyl-imidacarbocyanine iodide (JC-1), which selectively enter into the mitochondria, formed monomers (green color), indicative of depolarization into the mitochondrial membrane potential (ΔΨm) (remain as multimer J-aggregates (red color) in intact mitochondria). ΔΨm was analyzed by immunofluoresnce microscopy, (**E**) ΔΨm was evaluated by flow cytometry and percentile values of ΔΨm induced by *P. zopfii* in bMECs. \**P* < 0.05, \*\**P* < 0.01.

Transcriptional expression of genes regulating mitochondrion-mediated apoptosis, including increased *Bax* and *Apaf-1* (Supplementary Fig 1G-H) and decreased *Bcl-2*, were detected in bovine MECs inoculated with *P. zopfii* GT-II (Fig 5A). Apoptotic proteins, cytochrome-c and caspase-9 at early points (4 hpi) followed by caspase-3 later (24 hpi), increased after *P. zopfii* GT-II infection (Fig 5B-C). Likewise, cytochrome-c and cleaved caspase-9 and-3 were over time increasingly immune blotted (Fig 5D) and immunolocalized (Fig 5E) in bMECs infected with *P. zopfii* GT-II. Apart from decreased *Bcl-2* expression after 24 hpi, no effect of *P. zopfii* GT-I on apoptotic genes in bMECs was observed (Fig 5A).

**Fig 5.**
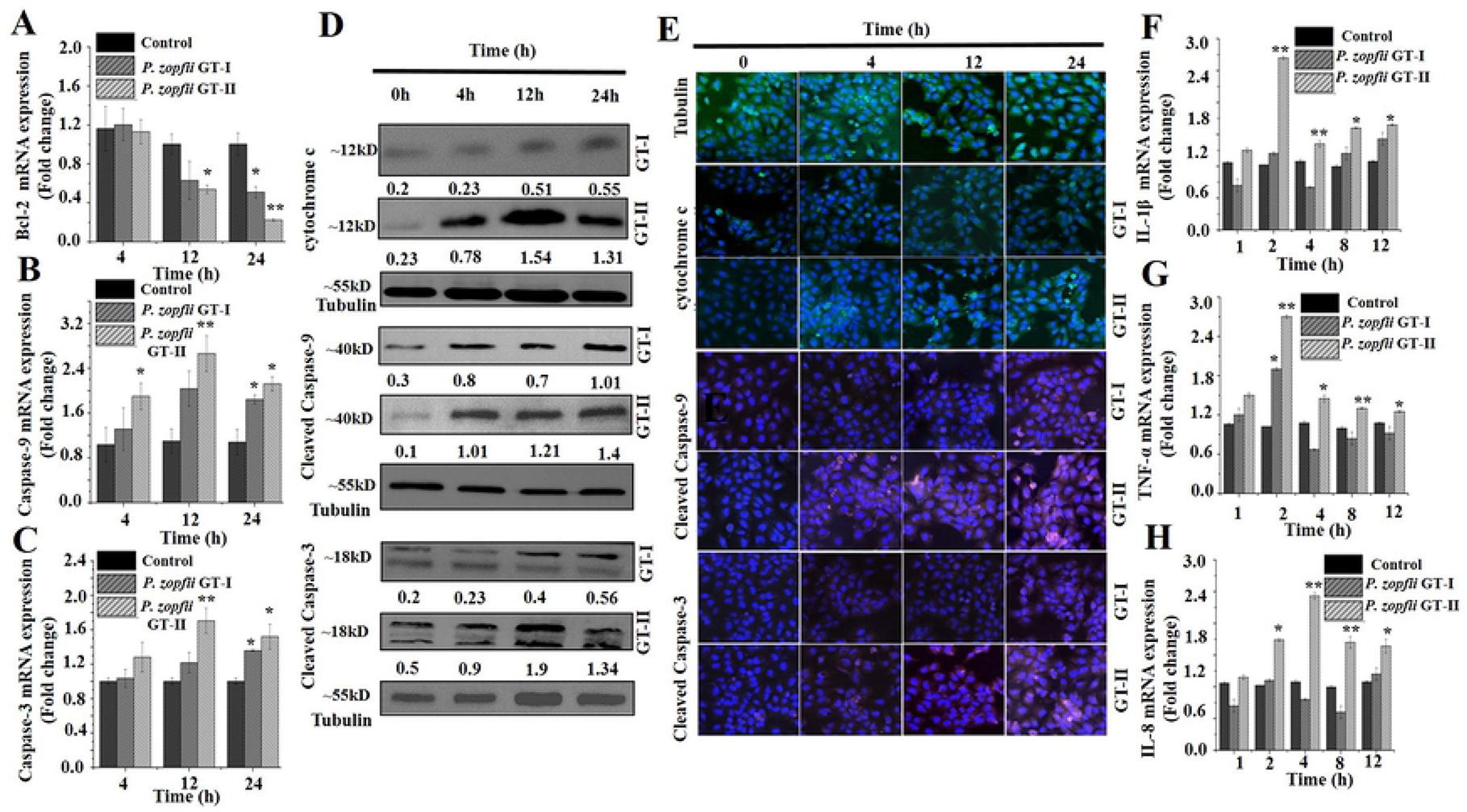
(**A, B and C**) Transcriptomic analysis of *Bcl-2*, caspase-9 and caspase-3, respectively. (**D**) & (**E**) Western blot and confocal laser scanning microscopic analysis of cytochrome-c, caspase-9 and caspase-3 in bMECs. (**F, G and H**) mRNA expression of pro-inflammatory cytokines (*TNF-α, IL-1β* and *IL-8*) quantified by qPCR in bMECs after infection of *Prototheca zopfii* genotype (GT)-I and -II infection. \**P* < 0.05, \*\**P* < 0.01.

Infection of *P. zopfii* GT-II induced pro-inflammatory responses in bMECs as demonstrated by upregulated mRNA expression of *IL-1β, TNF-α* and *IL-8* (after 2 hpi; Fig 5F-H). However, GT-I did not modify any inflammatory cytokine response in bMECs. Taken together, *P. zopfii* GT-II was demonstrated to cause udder disease by provoking apoptosis and inducing inflammatory cytokine expression in mammary epithelium.

## Discussion

Previously, pathogenesis of protothecal mastitis and virulence of *P. zopfii* GT-II isolated from bovine milk were uncertain. In this study, we described the pathogenic role of *P. zopfii* GT-II when initiating acute mastitis and mitochondrion-mediated apoptosis using a murine mastitis model and cultured mammary epithelial cells and macrophages. Our study demonstrated that *P. zopfii* GT-II invaded mammary parenchyma and caused acute mastitis, with severe infiltration of macrophages and neutrophils and marked epithelial damage. A destructive role of *P. zopfii* GT-II has been previously reported in the udder interstitium of cows and mammary acini of mice experimentally infected with *P. zopfii* GT-II [28,29].

Mammary epithelial cells are essential in microbial infection for sensing pathogens and producing an array of inflammatory cytokines [30]. Pro-inflammatory cytokines, including *TNF-α*, *IL-1β, IL-6* and *IL-8*, have a direct cytopathic effect leading to tissue damage [31]. Additionally, *IL-1β* and *TNF-α* can induce cell apoptosis [32]. *Prototheca zopfii* GT-II infection triggered expression of *IL-1β, Cxcl-1/IL-8*, and *TNF-α* in murine macrophages and bMECs. Thus, *P. zopfii* GT-II provoked apoptosis of bMECs by inducing *IL-1β* and *TNF-α* release in macrophages and mammary epithelial cells. *Prototheca zopfii* GT-II was more pathogenic than *P. zopfii* GT-I, commonly isolated as an enviromental apathogenic microbe. *Prototheca zopfii* GT-II induced more *IL-8* mRNA in bMECs compared to GT-I-inoculated or uninfected cells. Increased levels of *IL-8* mRNA in murine MECs and bovine MECs induced by *P. zopfii* GT-II demonstrated that mammary epithelial cells are an important source of *IL-8* and that this chemokine is key during protothecal mastitis, perhaps by recruiting leukocytes, as demonstrated by its chemoattractant role in *Staphylococcus aureus* infection in bMECs [33,34].

Whereas *P. zopfii* has been reported to induce apoptosis in cultured bMECs [20,21], we demonstrated the pro-apoptotic role of *P. zopfii* GT-II in a murine mastitis model. The pro-apoptotic character of *P. zopfii* GT-II was demonstrated by increased numbers of TUNEL-positive cells in *P. zopfii* GT-II-infected mice, along with reduced *Bcl-2* levels and elevated transcriptomic levels of *Bax*, *Apaf-1*, caspase-3, and caspase-9. These all indicated apoptosis via the intrinsic pathway, with functional alterations in mitochondria in mammary epithelial cells infected with *P. zopfii* GT-II. Moreover, *P. zopfii* GT-II induced ROS generation [21] which triggers mitochondrial *Bax*, a proapoptotic element of the *Bcl-2* family proteins [35]. *Prototheca zopfii* GT-II invaded bMECs and murine macrophages, and indeed, apoptotic effects were promoted by microbial internalization, but independent of phagocytosis. *Prototheca zopfii* GT-II had higher penetration capabilities in bMECs than *P. zopfii* GT-I. We propose that mitochondrial damage due to *P. zopfii* GT-II invasion released protein cytochrome-c from intermembrane spaces into cytosol, which bonded with *Apaf-1* to initiate apoptosome formation and activation of caspase-9 and caspase-3 [36–38]. Such *P. zopfii*-driven apoptosis was not restricted to mammary epithelial cells but also applied to leukocytes, including murine macrophages. A hypothetical schematic illustration of mitochondrial caspase-induced apoptotic pathway and NF-κB subunit 65 transiting into the nucleus in protothecal mastitis (Fig 6) was consistent with reports in bMECs, wherein *P. zopfii* GT-II regulated transcription of pro-inflammatory genes like *IL-1β* and *TNF-α* [33]. In conclusion, pathomorphological alteration caused by *P. zopfii* GT-II highlighted this gentoype as a mastitis pathogen capable of penetrating into mammary epithelial cells to induce inflammation and cell death, via mitochondrial-dependent apoptosis.

**Fig 6.**
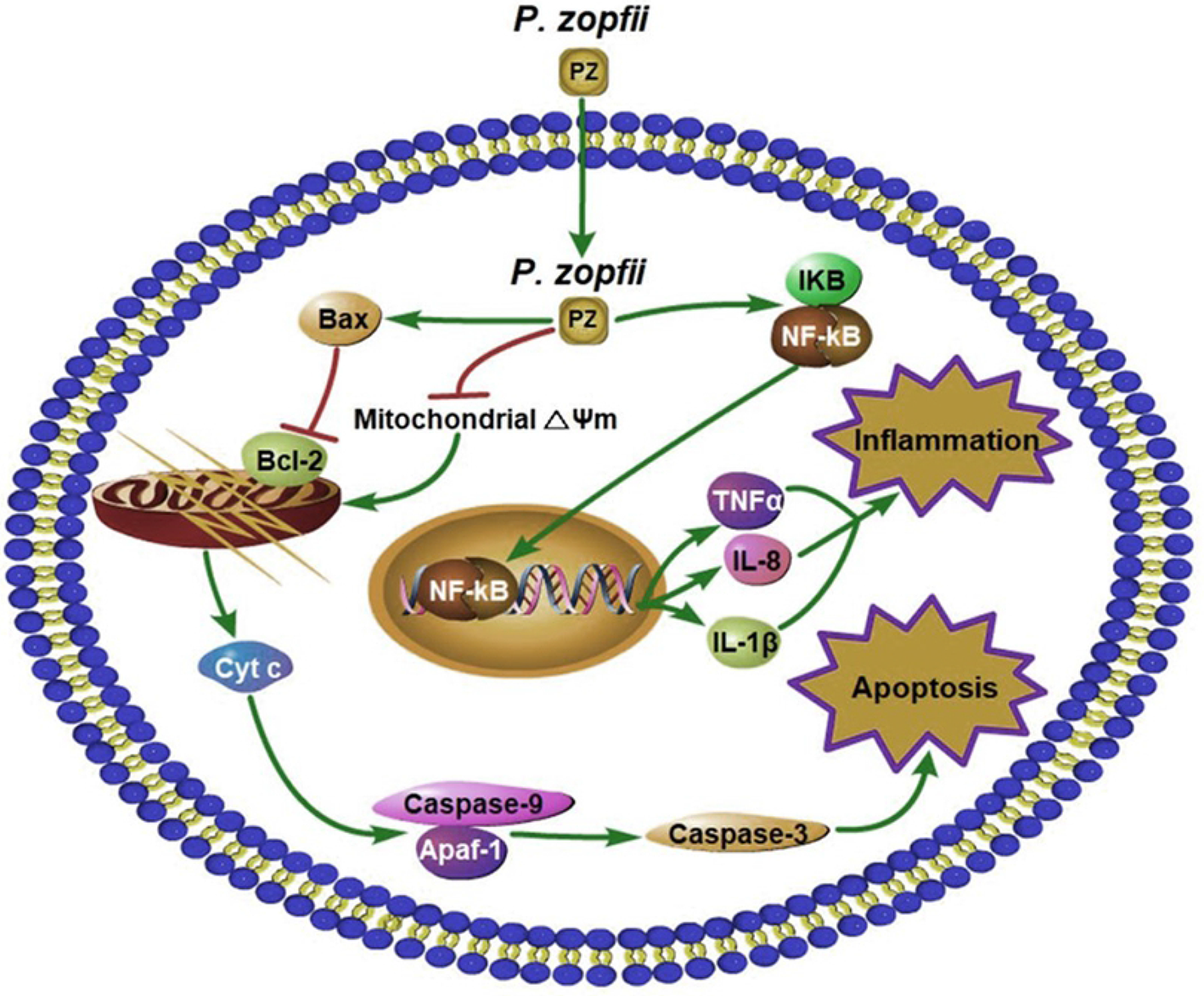
Schematic presentation of mitochondrial-caspase induced apoptosis and inflammation. Depolarization of mitochondrial transmembrane (ΔΨm) causes the release of cytochrome-c, which may initiate caspase cascade. Cytochrome-c bonds with apoptotic protease-activating factor 1 (*Apaf-1*) and activates caspase-9, this cleaves and activates caspase-3, which triggers apoptosis. NF-κB subunit 65 transiting into the nucleus wherein it regulates transcription of pro-inflammatory genes, e.g. *IL-1β* and *TNF-α*.

## Author contributions

MS, EC and HB conceived and designed the experiments. MS and PAC conducted animal sampling. MS, SX, and YL cultured and isolated *P. zopfii*. MS, JG and PAC prepared immunohistochemistry images for leukocytes and macrophage studies. MS, CK and PAC conducted the histopathological exams. MS, LC, JPK, CK and EC contributed to data analysis and interpretation as well as manuscript editing. MS, EC, HWB and HB drafted and wrote the manuscript.

## Acknowledgments

This study was financially supported by the National Natural Science Foundation of China (No. 31572587, 31772813 and 31850410474), the High-end Foreign Experts Recruitment Program (No. GDT20171100013), the National Dairy Industry and Technology System (CARS-37-02A), the Natural Sciences and Engineering Research Council (NSERC) Discovery Grant (RGPAS-2017-507827), the Eyes High International Collaborative Grant for Young Researchers (10014539) (University of Calgary) and Dairy Research Cluster 3 (1049122) for EC.

---

**Supplementary Fig 1.** mRNA expression of *Bax* and *Apaf-1* after infection of *Prototheca zopfii* genotype GT-II infection in mouse mammary tissues (**A and B**), murine MECs (**C and D**) and murine macrophages (**E and F**), whereas in bMECs (**G and H**) mRNA quantified after infection of *Prototheca zopfii* genotype (GT)-I and -II infection by qPCR and expressed as fold change relative to uninfected samples. \**P* < 0.05, \*\**P* < 0.01.

